# Stock identification of Mediterranean horse mackerel (*Trachurus mediterraneus*) through the analysis of morphometric characters in the Adriatic Sea

**DOI:** 10.1101/2024.07.23.604807

**Authors:** Claudio Vasapollo

## Abstract

Phenotypical differentiation among stocks of Mediterranean horse mackerel *Trachurus mediterraneus* in the Adriatic Sea was investigated through the analysis of several individual morphometric characters. Overall, 426 individuals of Mediterranean horse mackerels were sampled from the northern, central and southern Adriatic Sea in 2012 and 2013, and then compared using multivariate techniques (linear discriminant analysis). Discriminant analysis suggested a scarce intermingling between the northern and central Adriatic with the southern part. The former two areas showed more stable populations, overlapping both in space and time. The latter seemed to be more variable over the years. A possible hypothesis for this, to be investigated, is the flow of individuals from the Ionian and Aegean Seas populations through the Otranto Channel. When the morphometric variables were investigated, the results showed that most of the variability was associated with the head characters. In particular, the northern and central Adriatic Sea individuals had shorter and thicker heads than the southern ones. This could be due to different feeding habits: the former mainly feed on small fishes, the latter mainly on euphausiids. A short mouth could reduce the power of suction of bigger preys, while a long mouth could increase the volume of water to be filtered to feed on small planktonic crustaceans. From this study, it comes clear that the Mediterranean horse mackerel populations should not be managed as a single stock in the Adriatic Sea as it was evident that morphological two different populations are present in the basin.

## Introduction

Mediterranean horse mackerel *Trachurus mediterraneus* (Steindachner 1868) is a Carangiformes semipelagic species widely distributed in the Mediterranean and the Black Sea (Fischer, Bauchot & Schneider, 1987). Although this species, and its congeneric ones, represent an important resource in the rest of the Mediterranean Sea (Turan, 2004), no reliable landing data are available for the Adriatic Sea (Jardas, Santic & Pallaoro, 2004). Estimates of the mean annual landing values for the period 1970 – 2021 extracted from the GFCM - FAO (General Fisheries Commission for the Mediterranean – Food and Agricultural Organization of the United Nations) capture production database (www.fao.org/gfcm/data/capture-production/ar/), assessed the landings of *Trachurus* spp. (generally, the congeneric species are difficult to be recognized by fishers since the species are quite similar) in about 5089 metric tons/year for the whole Adriatic Sea, showing a tendency of reduction in catches starting from the middle ‘80s. Most of these catches (more than 80%) come from the Italian fisheries that alone reported a mean annual landing value of about 4274 metric tons year^-1^ (corresponding to approximately 1.5% of the total fisheries Italian catches). Economically, catches of *Trachurus* spp. from the Italian coasts for the period 2003 – 2020 have returned a mean profit of approximately 1.5 million euros/year (less than 1% of the total gain of the Italian fisheries; data from the Italian National Institute of Statistic [ISTAT]; www.istat.it). Notwithstanding the lower impact of the catches of this species in the Adriatic Sea, its ecological importance is not negligible, as it represents an intermediate between lower levels (plankton to anchovies and sardines) and top predators (jackfish and tunas) in the trophic web like any other small pelagic fish (Fréon et al., 2005). Following these considerations, scarce are the paper focusing on the biology of this species in the Adriatic Sea (Arneri, 1983; Arneri & Tangerini, 1984; Viette, Giulianini & Ferrero, 1997; Šantić, Jardas & Pallaoro, 2003a; Jardas, Santic & Pallaoro, 2004; Šantić, Radja & Paladin, 2011; Pešić et al., 2012; Palermino et al., 2021). Recognizing the extension of the fish stocks and their structures is essential to optimize fishery management and their yield (Begg, Friedland & Pearce, 1999). A misrecognition of the structure of exploited stocks of a species can potentially lead to overfishing and depletion of less productive stocks when multiple stocks are differentially exploited (Begg & Waldman, 1999). Moreover, the degree of exchange between stock members has remained a challenge to fisheries scientists and managers (Campana & Casselman, 1993; Begg, Friedland & Pearce, 1999).

The study of the morphometric characteristics is one of the well proven methods to identify different fish stocks living in different environmental conditions (Begg & Waldman, 1999; Begg, Friedland & Pearce, 1999; Cadrin, 2000). Contrary to the genetic analysis that proves evolutionary differences between stocks, the study of the phenotypic characters is useful to identify short-term environmental induced differences between stocks (Begg, Friedland & Pearce, 1999). This approach worked well in discriminating *Trachurus* spp. stocks in the Black Sea, in the Marmara Sea and in the Atlantic regions. For example, Murta (2000) found that *T. trachurus* (Linnaeus 1758) along the Iberian coasts formed different stocks from the Atlantic coasts of Morocco. Successively, in a wider study comprising northern Atlantic and Mediterranean Sea, Murta, Pinto & Abaunza (2008) showed six different stocks of *T. trachurus*, three of which were in the Mediterranean Sea.

Turan (2004) was able to identify at least three stocks of *T. mediterraneus* between Black Sea, the Marmara Sea and the Aegean and eastern Mediterranean Sea. Examples are available also for other small pelagic species worldwide such as *Scomber australasicus* (Cuvier 1832), *Engraulis encrasicolus* (Linnaeus 1758), *Scomber japonicus* (Houttuyn 1782) and *Sardina pilchardus* (Walbaum 1792) (e.g., Tzeng, 2004; Erdoğan, Turan & Koc, 2009; Erguden et al., 2009; Baibai et al., 2012).

Therefore, the purpose of this manuscript is to assess the existence of different stocks of Mediterranean horse mackerel in the Adriatic Sea, a semi-enclosed highly productive basin of the central Mediterranean Sea, characterized by different hydrodynamic features between northern, central and southern basin and several promontories and archipelagos that likely act as physical barriers to the expansion of the populations of this species and/or to the mixing of different populations.

## Material and Methods

A total of 426 individuals of *Trachurus mediterraneus* were collected along all the Adriatic Sea (Figure 1).

**Figure 1:**
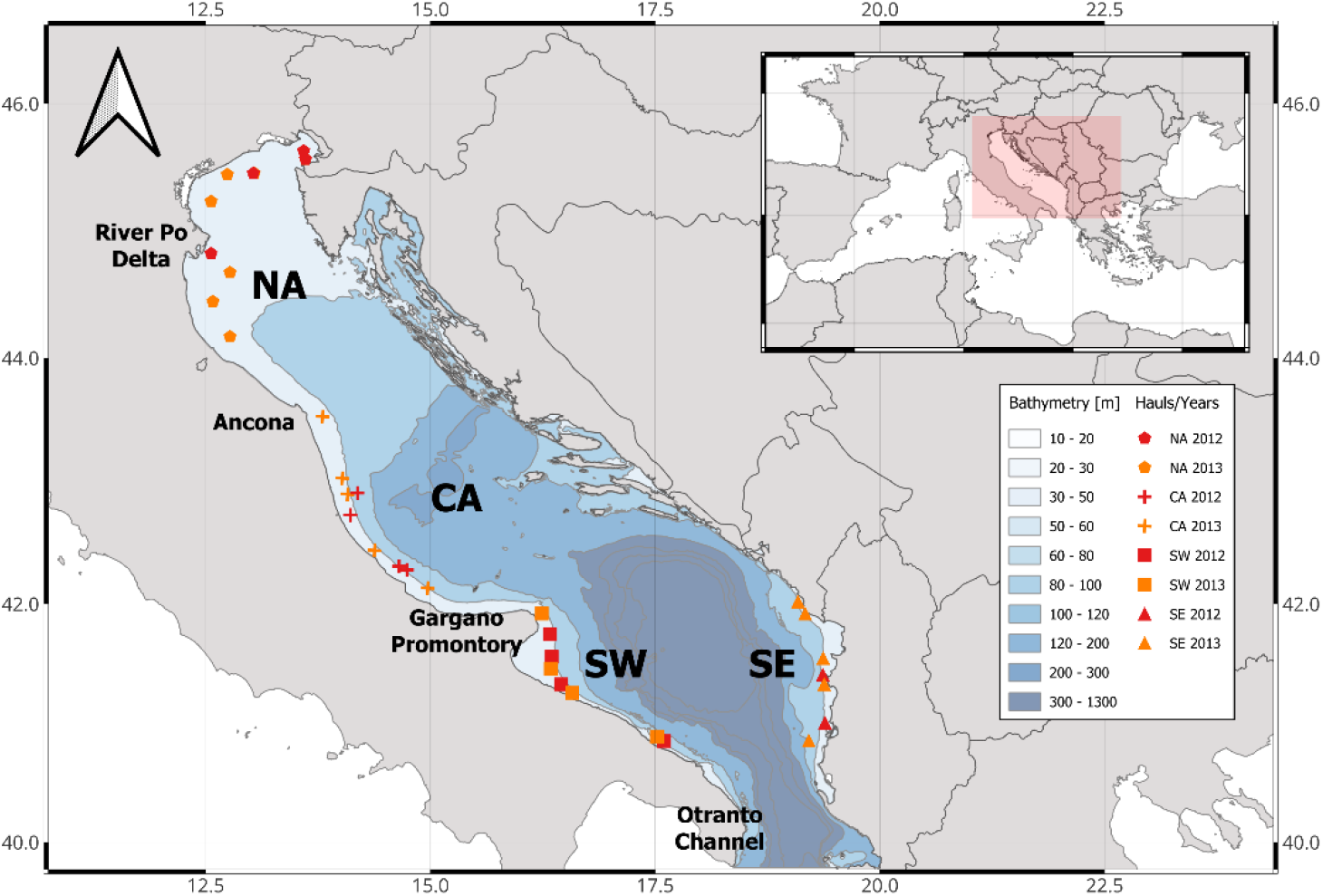
Map of the Adriatic Sea where specimen of *T. mediterraneus* were caught. The Adriatic Sea was divided into three main areas based on oceanographic characteristics: NA = North Adriatic; CA = Central Adriatic; SW = South-western Adriatic Sea; SE = South-eastern Adriatic Sea.

The available data on landings shows that *Trachurus* spp. yield in the whole Adriatic Sea declined between 1970 and 2020 (GFCM-FAO database [Figure 2], but see also Angelini et al. [2021]). On the Italian side of the Adriatic Sea, landing data clearly show this tendency at least for the period 2005 – 2020 (ISTAT; Figure 2) where it is evident that most of the landings came from the south of Italy (in particular, from Apulia region).

**Figure 2:**
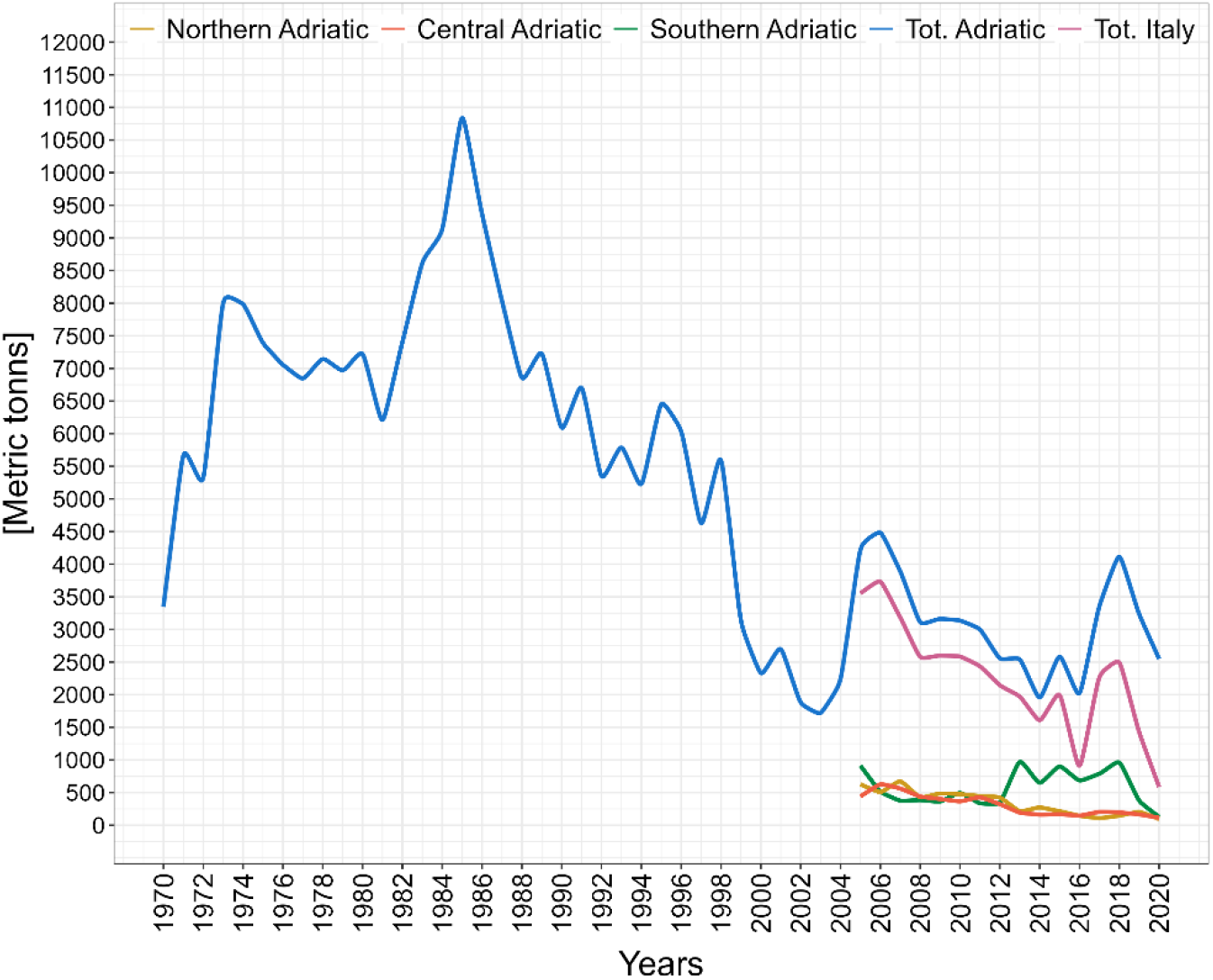
Trend of the total landings of *Trachurus* spp. in the Adriatic Sea and the rest of the Mediterranean Sea (Northern, Central, Southern and Total Italy data from the Italian National Institute of Statistics [ISTAT]; Total Adriatic [both western and eastern) from General Fisheries Commission for the Mediterranean – Food and Agricultural Organization of the United Nations [GFCM – FAO]).

The basin was ideally divided into 3 main areas (according to the mean cyclonic circulations and bathymetry; Artegiani et al., 1997): 1) Northern Adriatic (NA) from the Gulf of Trieste to the promontory of Ancona and with an average depth of 35 m; 2) Central Adriatic (CA) from the promontory of Ancona to the promontory of the Gargano mountain and an average depth of 140 m, with the two Pomo depressions reaching 260 m; 3) and southern Adriatic with an average depth > 500 m and a depression of > 1200 m, subdivided into South-Western (SW) and South-Eastern (SE) Adriatic. Specimen were obtained by means of pelagic trawls (Table 1) made during two echo-survey cruises held in 2012 and 2013 in the northern-central (September) and southern (July-August) Adriatic Sea during the reproductive period of this species (Arneri, 1983; Viette, Giulianini & Ferrero, 1997) in the framework of the MEDIAS project (MEDiterranean International Acoustic Survey; Leonori, Tičina & De Felice, 2011; Leonori et al., 2021).

**Table 1:**
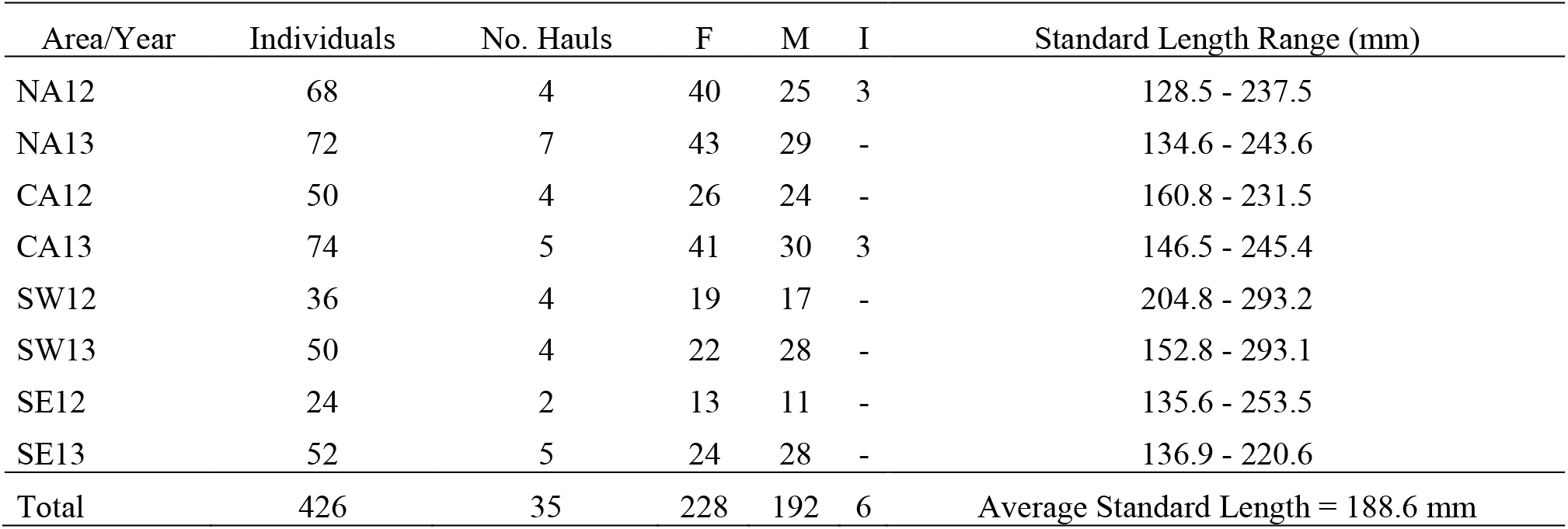
Areas of sampling, number of hauls from which specimen of *T. mediterraneus* come, number of individuals per area and year, number of females (F), males (M) and immature (I) individuals per area and range of the standard length per area. The grand mean of the standard length was 188.6 mm.

According to Reist’s (1985) recommendation that at least 25 individuals should be used for morphological analyses, a range of 24 to 68 individuals were collected in 2012 and 50 to 74 in 2013 (Table 1). The fish were kept deep-frozen (−20 °C) for transport to the laboratory, where several landmarks were pinned to the fishes before taking photographs (Canon PowerShot SX260 HS, 50 mm normal lens to avoid distortion of the image). By means of the image free software ImageJ v.1.54g (Abramoff, Magalhaes & Ram, 2004), a truss-network was built accounting for a total of 43 lengths, while three measures of width were taken by means of a digital caliper (Figure 3).

**Figure 3:**
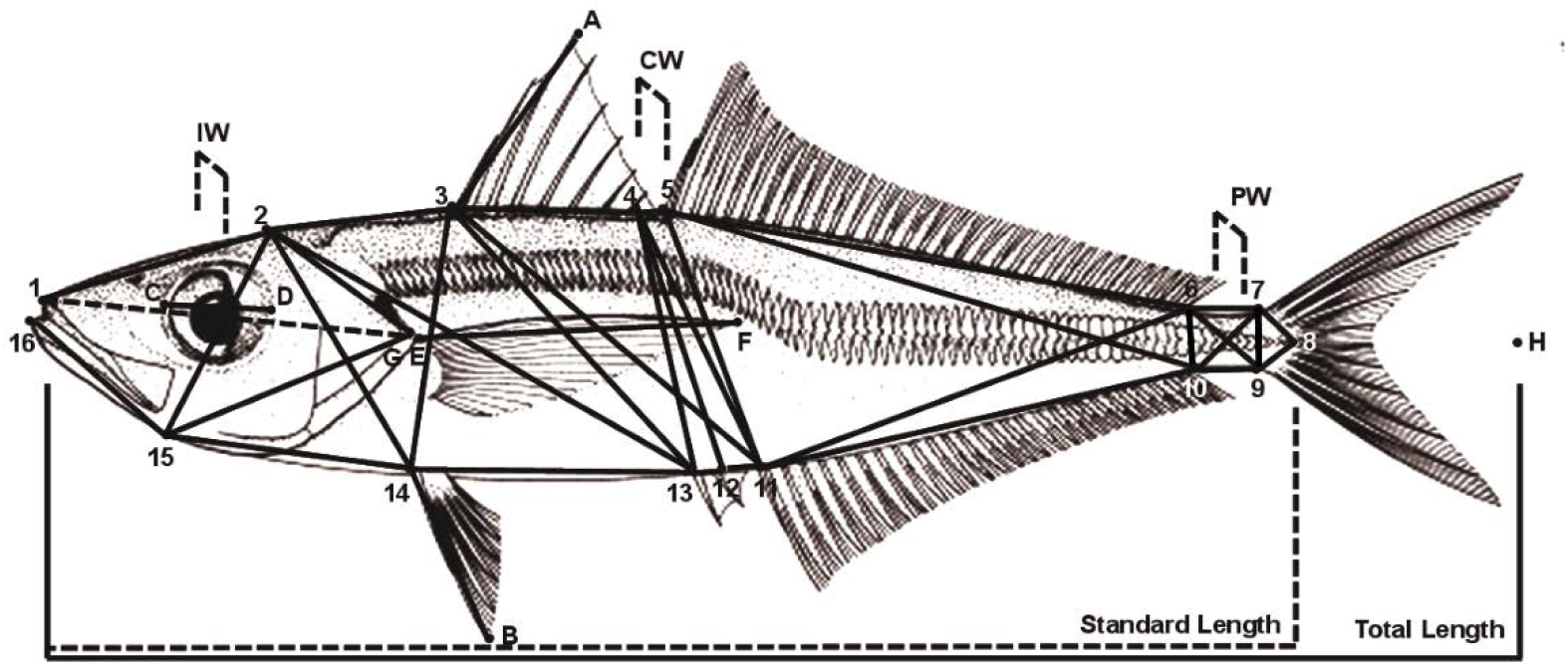
Locations of the 19 pins and 43 lengths of the truss network of *T. mediterraneus* used for the multivariate analysis. IW = interorbital width; CW = central width; PW = peduncle width; C_D = eye diameter; 1_G = head length; E_F = pectoral fin length; 14_B = ventral fin length; 3_A = first dorsal fin height.

To remove the size dependent effect of each morphometric measure the Elliott, Haskard & Koslow (1985) formula was applied:

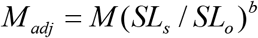

where M is the original morphometric measurement, *M*_*adj*_ the size adjusted measurement, *SL*_*s*_ the overall mean of standard length for all fish from all samples (188.6 mm) and *SL*_*o*_ the standard length of each single individual. The parameter *b* was estimated for each character as the slope of the regression of log *M vs* log *SL*_*o*_. Correlation coefficients between transformed measurements and standard length were calculated to check if the data transformation was effective in removing the effect size. If any correlation among variables persisted after transformation, one of the variables was maintained as a proxy for the others. The effect of sex on the measurements was not tested since Mediterranean horse mackerel do not show any sexual dimorphism (Jardas, Santic & Pallaoro, 2004; Turan, 2004).

After data standardization, an un-weighted pair-group method using arithmetic averages (UPGMA) cluster analysis of the Euclidean distances of the eight groups (eight areas) of individuals was performed to assess the presence of clusters of fish aggregations (Legendre & Legendre, 2012; Borcard, Gillet & Legendre, 2018). Once obtained the clusters, a Linear Discriminant Analysis (LDA) on standardized data was performed to identify the combination of variables that best separated *T. mediterraneus* samples. It proceeds in two steps: 1) one tests for differences in the morphometric variables among the predefined groups; 2) if the test supports the alternative hypothesis of significant differences among groups, the analysis proceeds to find the linear combinations (called discriminant functions) of the variables (standardized coefficients) that best discriminate among groups (Legendre & Legendre, 2012). After cross-validation (jackknife), each fish individual is allocated to the group with the nearest centroid, and the proportion of individuals allocated to each group is calculated after production of a confusion matrix. The proportion of correct allocation is taken as a measure of the integrity of that group. To test the assumption of multivariate homogeneity of within-group covariance matrices a permutational test has been applied (Borcard, Gillet & Legendre, 2018). To estimate the significance of the differences among groups, a Permutational Multivariate Analysis of Variance (PERMANOVA; Anderson, 2001) was performed between each pair of groups of a Euclidean matrix built with the Elliot’s transformed morphometric measurements.

Statistical analyses were performed with R packages stats v.4.4 (*hclust* function; R Core Team, 2024), MASS v.7.3 (*lda* function; Venables & Ripley, 2002) and vegan v.2.6(*adonis2, betadisper, permutest* functions; Oksanen et al., 2024).

## Results

A total of 426 individuals (average standard length = 188.6 cm; range size from 128.5 cm to 293.2 cm) have been collected for morphometric analysis during 2012 and 2013 cruises in the Adriatic Sea from a total of 35 hauls (Table 1). Most of the morphometric variables measured were highly size dependent but the Elliot’s transformation applied was effective for almost all the variables (Table 2). Notwithstanding, seven variables (1_8, 8_9, 4_12, 4_11, 5_10, 11_6 and 3_13) still showed multicollinearity, thus, to reduce their redundant effect on the analysis they were dropped from the list when the correlations were > 0.90 (Table 2), overall reducing the number of morphometric variables to 36.

**Table 2:**
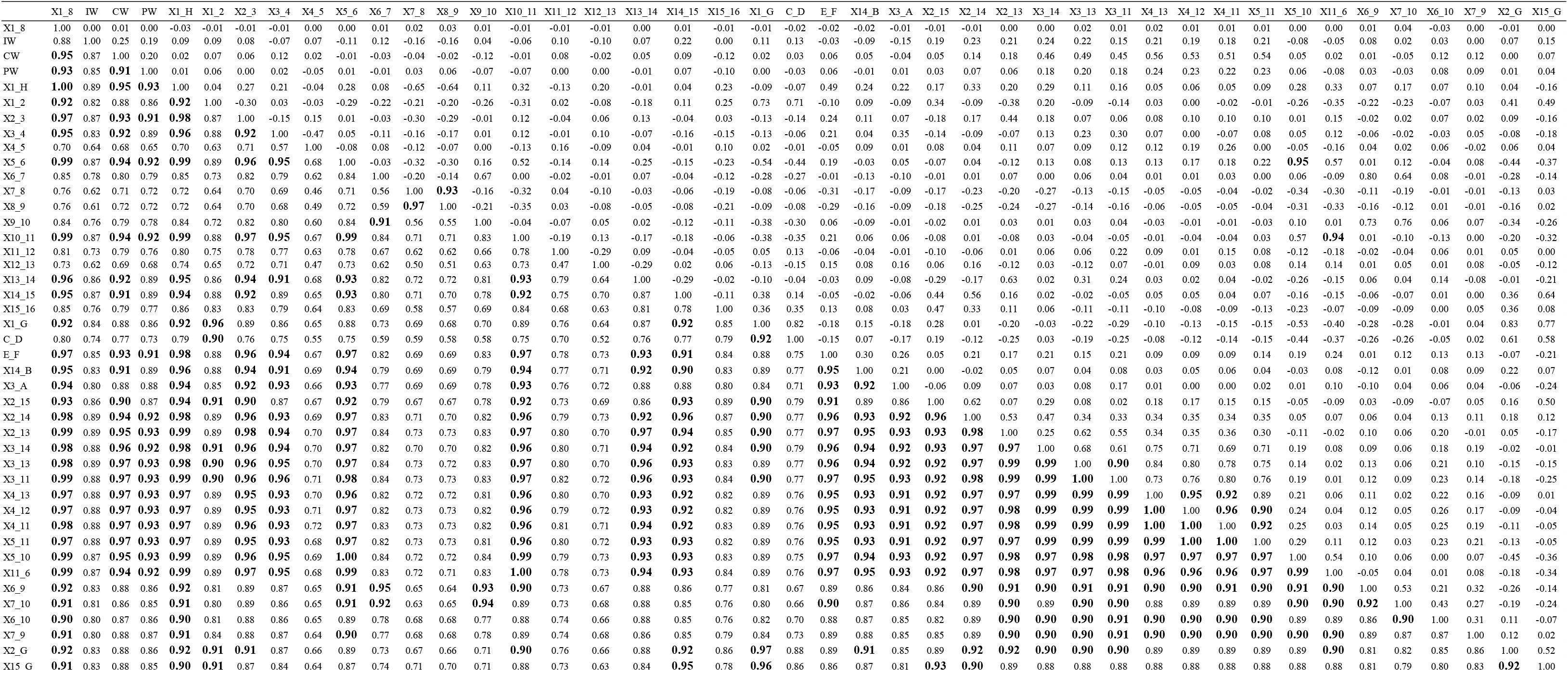
Correlation coefficients between characters, before and after the removal of the size effect, are respectively shown below and above the diagonal. Correlation values higher or equal to 0.90 are in bold.

The cluster analysis showed the division of individuals in mainly 2 groups based on the geographical separation between south and north-central Adriatic (Figure 4).

**Figure 4:**
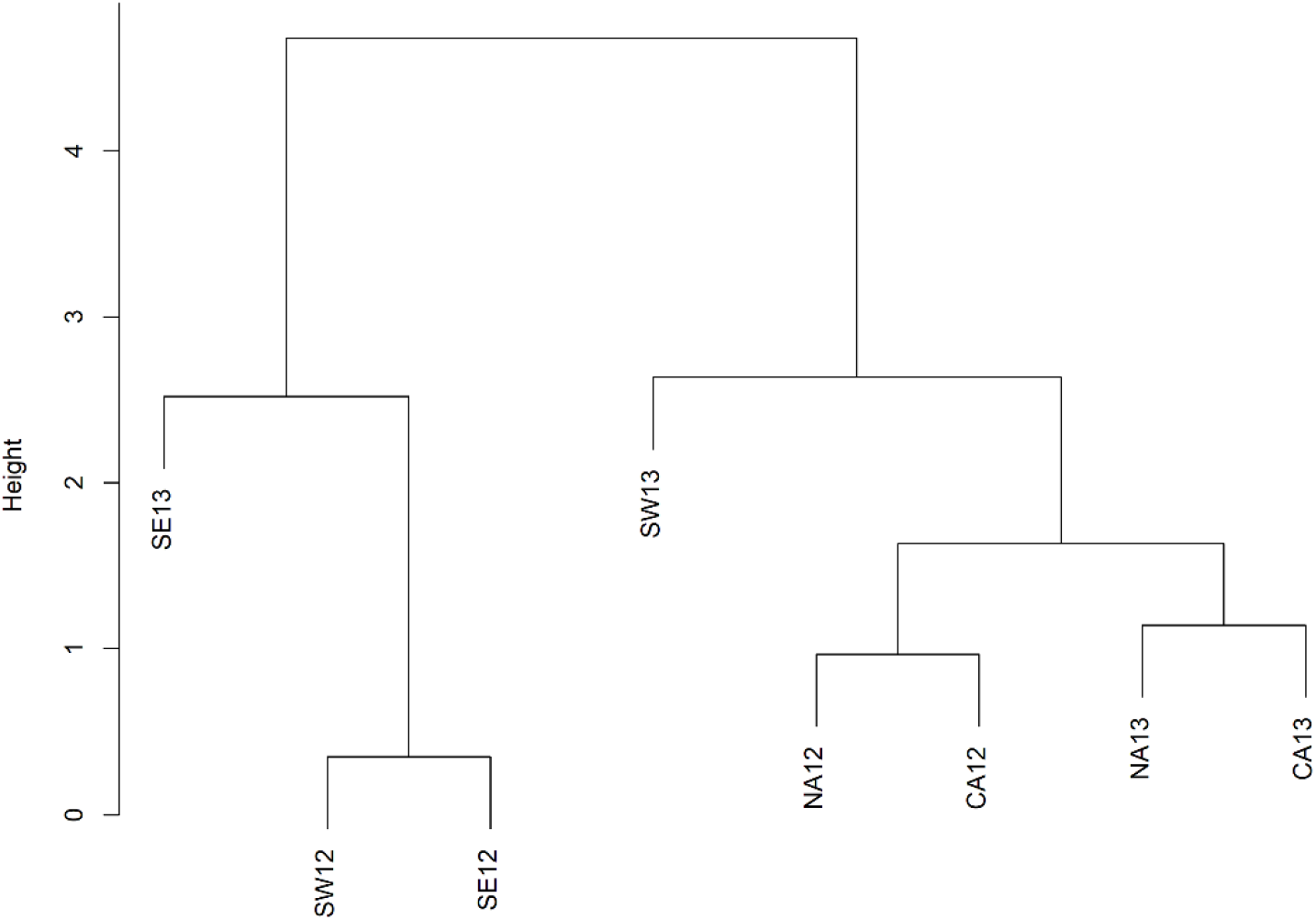
Dendrogram of the cluster analysis (UPGMA) for the centroids of the groups based on Euclidean distances of morphometric data. Cophenetic correlation = 0.93. NA = North Adriatic; CA = Central Adriatic; SW = South-western Adriatic Sea; SE = South-eastern Adriatic Sea.

Based on the cluster analysis, and following the hypothesis of the presence of different horse mackerel populations in the basin, the within-group analysis did not show any difference (Table 3). On the other hand, PERMANOVA clearly showed a difference between areas (Table 3), confirming the separation of the populations in different areas of the Adriatic Sea.

**Table 3:** Results of the PERMUTEST, to test the within-group multivariate homogeneity, and of the PERMANOVA, to test the between-groups differences. The number of permutations for both test was set to 9999.

| PERMUTEST |  |  |  | PERMANOVA |  |  |  |
| --- | --- | --- | --- | --- | --- | --- | --- |
|  | df | F | p |  | df | F | p |
| Area | 7 | 1.3 | 0.2667 | Area | 7 | 7.1 | 0.0001 |
| Residuals | 418 |  |  | Residuals | 418 |  |  |
|  |  |  |  | Total | 425 |  |  |
| No. Perm = 9999 |  |  |  | No. Perm = 9999 |  |  |  |

The LDA analysis plot showed a clear separation between the northern/central areas respect to the southern area, suggesting the existence of at least two populations of Mediterranean horse mackerel (Figure 5). Moreover, the northern/central area groups maintained a temporal stability between the two years being all groups close each other, while, on the other hand, the southern area groups were scattered showing temporal instability and suggesting the further presence of ephemeral populations changing each year.

**Figure 5:**
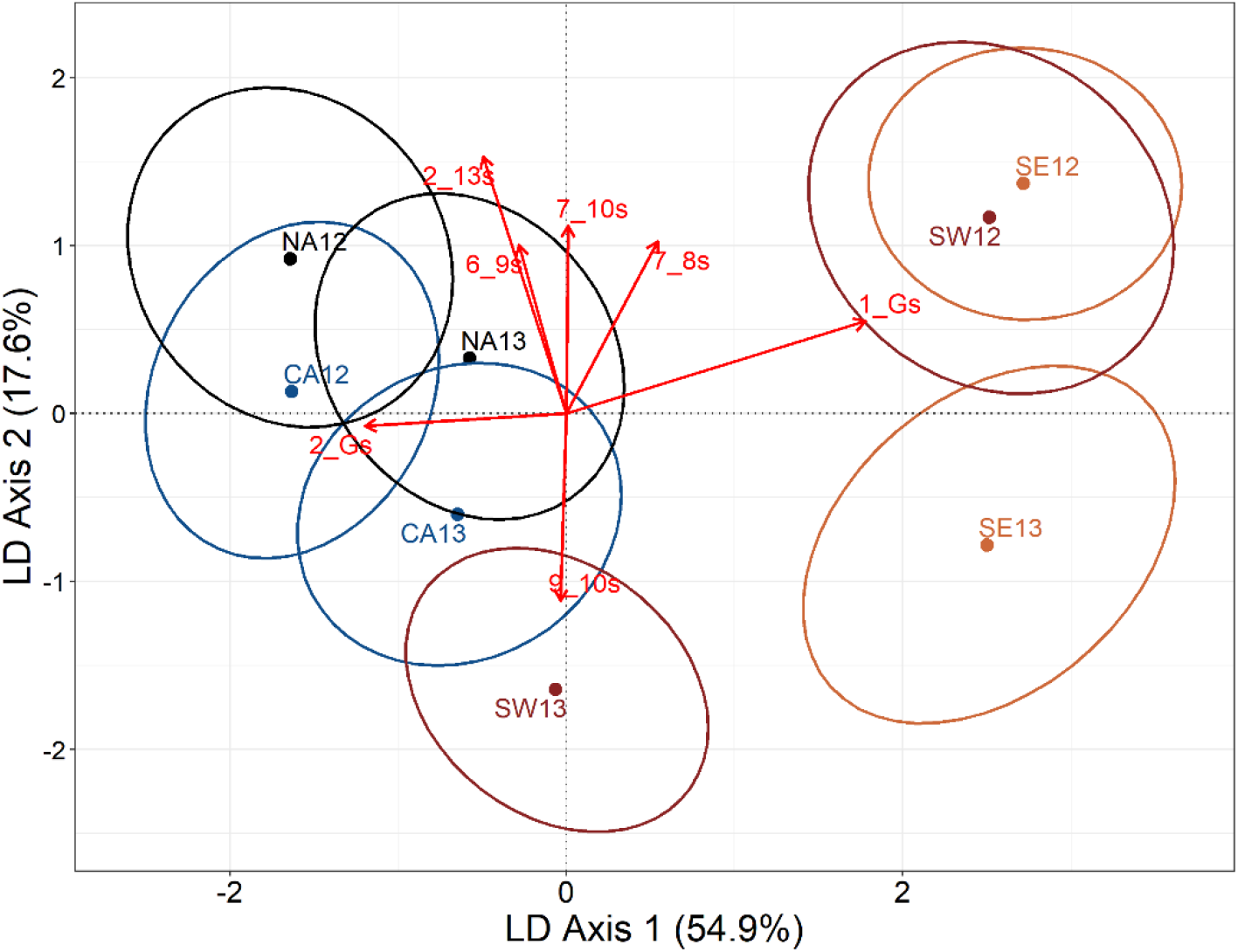
Plot of the centroids of the sample groups in the space defined by the Linear Discriminant Analysis. Both axes explained together 72.5% of the total between-group variability (first axis 54.9%, second axis 17.6%). Dots represent centroids of the samples per area per year. The radius of the ellipses around each centroid corresponds to one standard deviation of the Euclidean distances from each individual to its group centroid. Arrows represent the contribution of morphometric variables to the canonical functions (standardized coefficients). Only the most correlated vectors (≥ |1|) are reported. NA = North Adriatic; CA = Central Adriatic; SW = South-western Adriatic Sea; SE = South-eastern Adriatic Sea.

The first two discriminant axis produced by LDA explained most of the total between-group variability in the dataset (72.5%; LD axis1 = 54.9% and LD axis 2 = 17.6%), showing clear between-groups differentiation. The confusion matrix produced by the discriminant analysis (Table 4) showed that the cross-validated percentages of perfect classification of the individuals into the corresponding groups was 51.4%. The higher value of good correspondence was in the area SE13 (67.3%) while the lowest value was in CA12 (34%).

**Table 4:** Confusion matrix produced by the Linear Discriminant analysis. Numbers (and percentages) represent *T. mediterraneus* individuals correctly classified into their original group, in the validation of the discriminant analysis for the morphometric data. Rows are the original sample group and columns the reallocation group after jackknife cross-validation. r-SUM = row sum; c-SUM = column sum.

|  | NA12 | NA13 | CA12 | CA13 | SW12 | SW13 | SE12 | SE13 | r-SUM |
| --- | --- | --- | --- | --- | --- | --- | --- | --- | --- |
| NA12 | 40(58.8%) | 7 | 13 | 2 | 0 | 4 | 1 | 1 | 68 |
| NA13 | 10 | 35(48.6%) | 4 | 12 | 2 | 7 | 1 | 1 | 72 |
| CA12 | 22 | 2 | 17(34%) | 8 | 0 | 1 | 0 | 0 | 50 |
| CA13 | 4 | 19 | 5 | 34(46%) | 1 | 11 | 0 | 0 | 74 |
| SW12 | 1 | 3 | 1 | 1 | 18(50%) | 0 | 6 | 6 | 36 |
| SW13 | 0 | 8 | 2 | 12 | 0 | 26(52%) | 0 | 2 | 50 |
| SE12 | 0 | 0 | 0 | 1 | 4 | 0 | 14(58.3%) | 5 | 24 |
| SE13 | 0 | 0 | 0 | 3 | 7 | 3 | 4 | 35(67.3%) | 52 |
| c-SUM | 77 | 74 | 42 | 73 | 32 | 52 | 26 | 50 | 426 |
| Percentage of Correct Classification = 51.4% |  |  |  |  |  |  |  |  |  |

To make the plot as much clear as possible, standardized coefficients were selected based on the absolute values (≥ |1|) to highlight those variables that mostly driven the spatial configuration (Figure 5). Two variables were strongly associated with the first most explanative axis corresponding to the length of the head and to the distance from the upper portion of the head until the operculum opening of the individuals (1_G and 2_G). On the other hand, some of the variables corresponding to the peduncle of the caudal fin and of the anterior part of the body were associated with the vertical axis (Figure 5).

## Discussion

Morphometric characters of Mediterranean horse mackerel showed significant phenotypic distinctness between north-central and southern Adriatic Sea. The present results suggested at least two different populations based on phenology, even if the southern groups are likely as different each other in both years as respect to the north-central population. Even though, a certain degree of mixing is plausible above all between the two populations at the border of the two areas CA and SW (at least for one year), but in any case, it seems not the rule but rather an exception, although more data are needed to support this statement. The individuals, from the north and the central Adriatic, were well mixed each other in both years. Under these circumstances, it is plausible to indicate the northern and central Adriatic populations as (at least morphologically) closed and self-recruiting populations because of their temporal stability. A clear idea of this hypothesis could potentially come from a further genetic study confirming (hopefully) the segregation of this population respect to the southern Adriatic and even from the rest of the Mediterranean. In fact, the southern individuals seemed to be quite different each other in both years. A possible explanation could be the influence of Mediterranean populations of *T. mediterraneus* entering the Adriatic Sea toward the Otranto Channel, thus favoring the constant flow into the southern Adriatic of new individuals each year. It is likely that the conditions (both environmental and feeding conditions) in the southern Adriatic are quite similar to the Ionian and Aegean Sea. In effect, the south Adriatic has open sea water mass characteristics as the northern Ionian Sea and the surface transport of water masses into the southern Adriatic occur from the eastern Ionian Sea (Vilibić et al., 2012) likely transporting individuals from Greek populations. This hypothesis should be proven by studying both southern Italian Ionian and Greek populations of Mediterranean horse mackerel both genetically and morphologically. The separation among populations might be due to the considerable differences in environmental conditions such as temperature, salinity and food availability between the three regions here considered (Artegiani et al., 1997). In fact, it is well known that fish exhibit morphological variation induced by the environment (Cadrin, 2000). Factor such as food availability (but also temperature and salinity among others, but even prolonged swimming; Turan, 2004) may determine phenotypic differentiation that, although not directly indicating gene flow among populations, may indicate that fishes spent their lives in separate regions. One of the clues that lead to the conclusion of individuals living in different habitats came from the discriminant analysis that showed that morphometric differentiation among samples was largely due to differences in the head features of fish. Also Turan (2004) found the same Mediterranean horse mackerel head differences along the Turkish coasts demonstrating that the most important leading variables in discriminating among stocks are associated with the prey selection in the different habitats. In fact, as reported by Gatz (1979), the relative head length would be linked to prey size. From the present results emerged that the northern and central Adriatic individuals were characterized by shorter but thicker head than the southern populations. This is not negligible since the diets between northern and southern populations are different enough to further support the distinction between Adriatic populations. Šantić, Jardas & Pallaoro (2003b) observed that the main preys of Mediterranean horse mackerel in the Adriatic Sea were plankton euphausiids, representing the 50% of the preys found in the stomachs analyzed. The rest of the food was represented by teleosts (as secondary food) and other planktonic crustaceans (as occasional food).

Euphausiisds are representative of deep and open waters, and, in fact, they are present in the southern and deepest parts of central Adriatic while in the north this planktonic group is not present (Šantić, Jardas & Pallaoro, 2003b). It is true that the prey size preference changes with fish size class (individuals under 28 cm total length prefer the planktonic crustaceans than small teleosts; Šantić, Jardas & Pallaoro (2003b)), but the size class ranges here analyzed are comparable among the areas and years, thus the kind of diets are almost all represented and the differences found between the head lengths corresponded to effective phenotypical differences due to different habitats. So, the differences in the head measures could be interpreted with respect to the kind of prey a Trachurus population feed on. For example, the northern and central populations potentially feeding mainly in small teleosts may be advantaged by having a short and thick mouth to facilitate the suction of the preys, while on the contrary, a long mouth could facilitate filtering a large volume of water to feed on plankton euphausiids. Regarding the morphometric characteristics associated with the second discriminant axis, it is more difficult to explain them. Notwithstanding, it is recognized that the kind of feeding habits promote changes in the body morphology of fishes (Blake, 2004). For example, the measure 2_13 (corresponding to the 2_12 in Turan, 2004), as well as the peduncle measures, is likely correlated to the swimming optimal functional design of the’s fish body. To maximize thrust and minimize drag a streamlined body is more effective, as those of the northern Adriatic individuals that need a rapid movement to catch fishes to feed on. The same solution is needed also to escape from predators (e.g., tunas), and this could explain why also individuals from the southern Adriatic (at least for the 2012) share these characteristics with the northern ones but could potentially due also to other intrinsic and/or extrinsic reasons. Interestingly, individuals from southern and central Adriatic caught in 2013, showed an opposite body shape respect the 2012 individuals. This should be investigated further. Nevertheless, these considerations should be viewed as speculative mostly because the second discriminant axis explained variance is quite low and thus, they should be taken with precaution.

## Conclusion

It is certain that for a reliable identification of several stocks a multi-methodological approach should be used (Begg & Waldman, 1999). The results of the present work should be compared and/or confirmed by analyzing data obtained from other phenotypic characteristics and genetic studies even from different sites (and possibly from the eastern Adriatic Sea). For example, another approach might be the study of otolith morphometry and chemistry that is increasingly used to discriminate between fish stocks (Campana & Casselman, 1993; Turan, 2006; Jemaa et al., 2015). So far, the Mediterranean horse mackerel in the Adriatic Sea is treated as a single stock (Angelini et al., 2021), but the present morphometric results showed that at least more than two stocks are present in the Adriatic waters. Unfortunately, no morphometric data are available from the northern-eastern and central-eastern Adriatic basin, and this is a lack for the present study, but already the results from the present study suggest that from a management point of view, the existence of different stocks could have huge consequences, above all if such population structure persists over time. If persistence would occur, it could be important to manage in different ways the different stocks because any depletion in one of them is unlikely to be compensated by immigration from other stocks, at least in a short time. Thus, for accurate assessment of the state of a stock, not only the boundaries but also the mixing levels between stocks should be considered (Turan, 2004; Murta, Pinto & Abaunza, 2008).

